# Memory-specific E-I balance supports diverse replay and mitigates catastrophic forgetting

**DOI:** 10.1101/2025.09.19.677474

**Authors:** Tomas Barta, Tomoki Fukai

## Abstract

The hippocampus is the brain’s central locus for memory processing. In a widely accepted hypothesis, the hippocampus stores short-term memories in attractor states, which are then consolidated in the neocortex as long-term memories. Hippocampal replay activities play a pivotal role in memory consolidation, but memories are only transiently, not persistently, replayed, casting doubt on the attractor memory hypothesis. Concepts like memory capacity, the upper bound on the number of storable stable memories in a neural network, may also need to be revised if their offline transient activation is more essential for memory consolidation. Here, we explore the biologically plausible mechanism of offline memory processing by asking a recurrent spiking network to replay reliably as many stored assemblies as possible. We demonstrate the importance of memory-assembly-specific inhibitory plasticity for replaying diverse memory assemblies. Furthermore, our results highlight the role of transient memory states in mitigating catastrophic forgetting.

## Introduction

The hippocampal area CA3 is thought to store memories in recurrent connections formed between excitatory pyramidal neurons through synaptic plasticity [1]. Theoretical studies have shown that recurrent neural networks can generate attractor dynamics going to persistent memory states, suggesting the attractor memory hypothesis. As recurrent connections generate crosstalk between attractors, the number of memory states storable in and retrievable from an attractor network is limited [2, 3]. A large body of theoretical studies exists on memory capacity in associative memory models of binary or rate-coding neurons [2–6], and modern network models emerging from machine learning research can give much stronger attractor states with a significantly large memory capacity [7, 8]. However, achieving a capacity as large as in rate-coding networks is challenging in spiking neural networks [9, 10]. Stabilization of stable memory states may be even harder in cortical circuits, as mechanisms such as spike frequency adaptation (SFA) are ubiquitous and can further destabilize such states[11–14].

It is widely believed that spontaneous replay of memory traces is essential for the consolidation of episodic memory during sleep [15–18]. If the hippocampus frequently replays an activity pattern experienced during an episode, the episode will be reliably stored in cortical memory storage. Conversely, if a memory trace is not replayed sufficiently often, the episode will not become a permanent memory. Memory-associated hippocampal assemblies are only transiently, but not persistently, reactivated during sleep. This observation challenges the conventional view that the hippocampus employs stable attractor states to memorize episodes. Rather, the hippocampus may transiently and repeatedly reactivate cell assemblies encoding episodes, and such reactivation can suffice for consolidating hippocampal memory traces into cortical storage.

Based on this hypothesis, here we construct recurrent neural networks of spiking neurons to clarify the dynamical properties and computational advantages of transient memory networks. In particular, we are interested in uncovering the inhibitory plasticity mechanism that maximally utilizes the transient memory states. Evidence is emerging for the importance of the plasticity of inhibitory neurons in stabilizing network activity [19–26]. The roles of inhibitory plasticity in memory storage have been studied in neural networks that embed a single or a small number of memory assemblies [27–30]. However, it remains largely unknown how inhibitory plasticity regulates the storage and retrieval performance of memory traces in recurrent neural networks.

To study the performance of transient memory networks at high memory load, we quantify the memory capacity of such networks based on the diversity of replay patterns during spontaneous activity. Replay diversity of transient memory states naturally extends the classic notion of storage capacity for stable memory states. The higher replay diversity, the more memories the networks possibly consolidate into long-term memory. Our results reveal that the details of inhibitory plasticity mechanisms are crucial for the performance of memory replay. The frequency and diversity of replayed memory assemblies, as well as their robustness against an increase in the number of embedded assemblies, depend crucially on the structure of the E-I balance organized by a specific inhibitory plasticity rule. Furthermore, the network can replay transient memory states even when the memory load is too large to sustain stable attractor states. Thus, this study suggests that transient memory states give a possible solution to catastrophic forgetting, the difficulty that stable memory states are abruptly destabilized when the number of memory assemblies exceeds a critical capacity.

## Results

### Global vs. local inhibitory plasticity rules

We modeled a network of 8000 excitatory and 2000 inhibitory spiking neurons equipped with a spike frequency adaptation (SFA) mechanism. We hard-wired a large number of memory assemblies in non-modifiable excitatory-to-excitatory (E-to-E) connections with log-normally distributed synaptic strengths. We made inhibitory-to-excitatory (I-to-E) connections plastic to stabilize network activity (Fig. 1A). Each memory assembly could be only transiently activated due to SFA. The assemblies were relatively sparse: each neuron had a 1% chance of belonging to any given assembly. Therefore, each assembly consisted of 80 neurons on average.

**Fig. 1.**
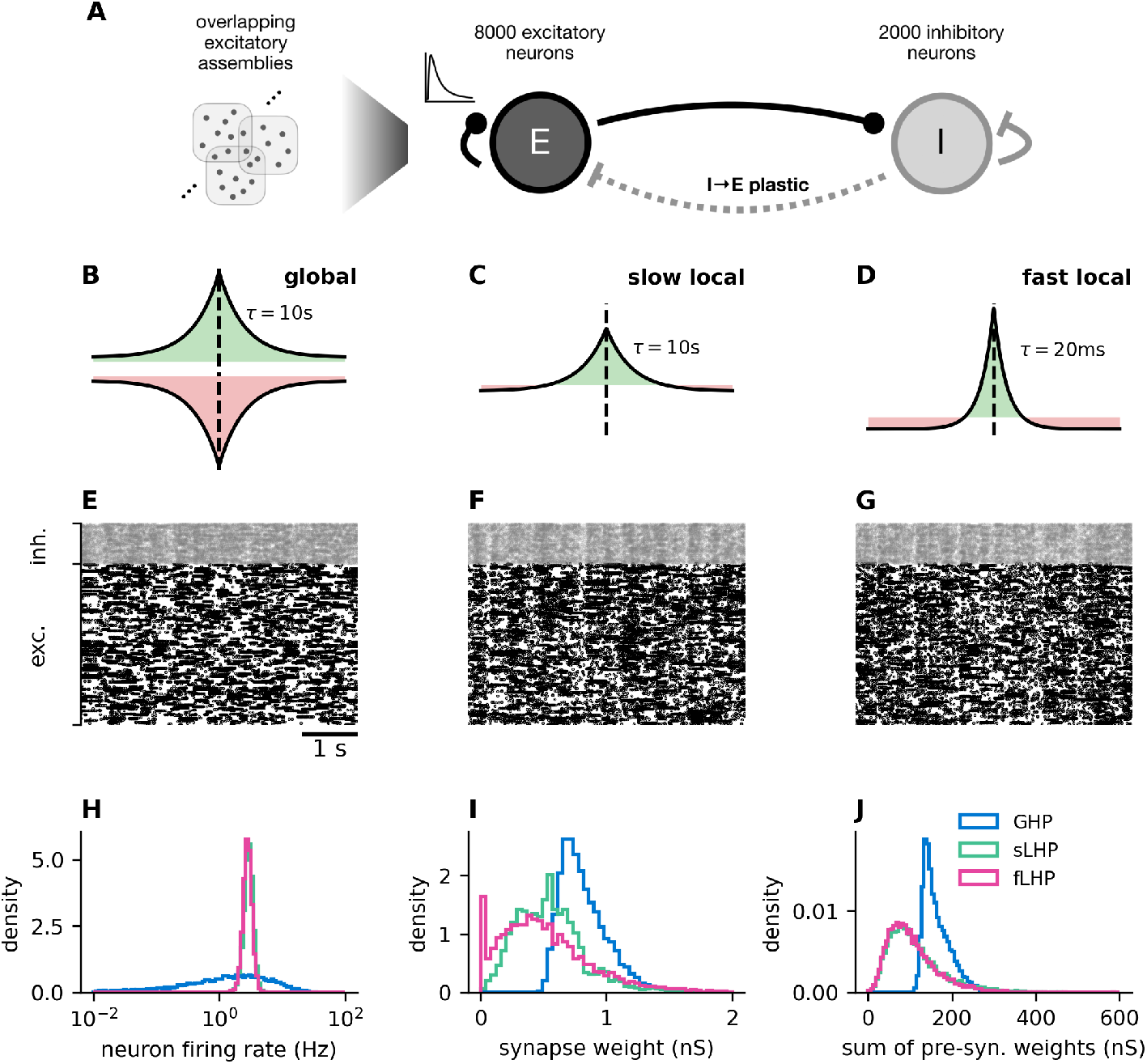
Inhibitory structures for different plasticity rules. **A:** The schematic illustration of the network model. **B-D:** Inhibitory plasticity rules. The GHP rule (B) strengthens (green) or weakens (pink) synaptic weights whenever the population firing rate is above or below threshold, respectively. The sLHP (C) and fLHP rules (D) are symmetric Hebbian rules. The two rules differ only in the decay time constant. **E-G:** Raster plots of spontaneous activity of the networks stabilized with the three rules (E: GHP, F: sLHP, G: fLHP). Spike trains of randomly chosen 800 excitatory and 200 inhibitory neurons are shown. **H:** Firing rate distributions are shown for the three rules. The distributions overlap almost completely for the sLHP and fLHP rules. **I:** The distributions of inhibitory synaptic weights over the entire networks are plotted. **J:** The distributions of inhibitory synaptic weights summed over individual excitatory neurons are plotted. The distributions highly overlap for the sLHP and fLHP rules.

As the biological rules of inhibitory plasticity remain elusive, we used three inhibitory plasticity rules to stabilize the activity of excitatory neurons. These rules may organize I-to-E connections with various levels of complexity:

1. *A network activity-dependent rule* stabilizes the mean firing rate of excitatory neurons [28] (global homeostatic plasticity, GHP rule; Fig. 1B).
2. *A neuron-specific rule* individually stabilizes the mean firing rate of each neuron on a slow timescale (slow local homeostatic plasticity; sLHP rule; Fig. 1C).
3. *A neuron-and-stimulus-specific rule* stabilizes the mean firing rate of each neuron, with inhibition tracking the excitation in time [27] (fast local homeostatic plasticity; fLHP rule; Fig. 1D).

The first rule induces potentiation or depression when the mean firing rate of all excitatory neurons is above or below a threshold, respectively, thus regulating the homeostasis at the whole network level [28, 29]. The second and third rules are symmetric Hebbian spike-timing-dependent plasticity (STDP) rules with a constant depression factor at each pre-synaptic spike [27]. The difference between the two rules is in the length of the STDP time constant. The slow time constant for sLHP (10 s) provides a blanket lateral inhibition, achieving local but loose EI balance, whereas the short time constant for fLHP (20 ms) allows strengthening of inhibitory synapses on the most active excitatory neurons, achieving tight local EI balance [31]. Below, LHP refers to both fLHP and sLHP. These local plasticity rules enable us to investigate the effects of a cell-assembly-specific excitation-inhibition (E-I) balance on memory processing with transient states. After training the network, we fixed the synaptic strengths to isolate the effect of the inhibitory structure from the potential effect of long-term synaptic plasticity on the network dynamics.

All three rules produced asynchronous activity with occasional spike bursts (Fig. 1E-G), a hallmark of networks with heavy-tailed synaptic connectivity [32]. Due to its global nature, the GHP rule preserved the heterogeneous firing rates of excitatory neurons, which were induced by a high variance in the total excitatory input to each neuron. The resultant distribution of firing rates seems consistent with highly skewed, typically lognormal, distributions observed ubiquitously in cortical networks [33, 34]. In contrast, both sLHP and fLHP rules produced a much more homogeneous distribution of excitatory firing rates around the target firing rate of postsynaptic neurons (Fig. 1H).

From GHP through sLHP to fLHP, increasingly finer tuning of the excitation-inhibition balance self-organized, leading to a broader and more complex I-to-E projection structure and higher variability in the I-to-E synaptic strengths (Fig. 1I). While the synaptic strength distributions displayed prominent differences between the sLHP and fLHP rules, they produced similar distributions of the sum of pre-synaptic inhibitory weights over the excitatory neuron population (Fig. 1J). This implies that the slow and fast rules generated different microscopic inhibitory circuit structures at the single-cell level but a similar macroscopic structure at the network level. Note that the finer inhibitory circuit structure enables a weaker total pre-synaptic inhibitory weight to stabilize network activity.

### The concept of memory capacity in spontaneous replay activity

Owing to SFA, the embedded memory assemblies formed transient attractors. Each network was capable of spontaneously and transiently replaying the embedded assemblies - an event during which all (or almost all) neurons of the assembly are active (Fig. 2A). Regardless of the self-organized inhibitory circuit structure, the replay frequency decreased rapidly with an increasing number of embedded memory assemblies, indicating weakening of the transient attractors (Fig. 2B). This is also evidenced by the decreasing average duration of each assembly activation (Fig. 2C). Additionally, transient assembly activations are, on average, the longest with the sLHP rule and shortest with the GHP rule, suggesting that the sLHP rule forms the strongest transient attractor states.

**Fig. 2.**
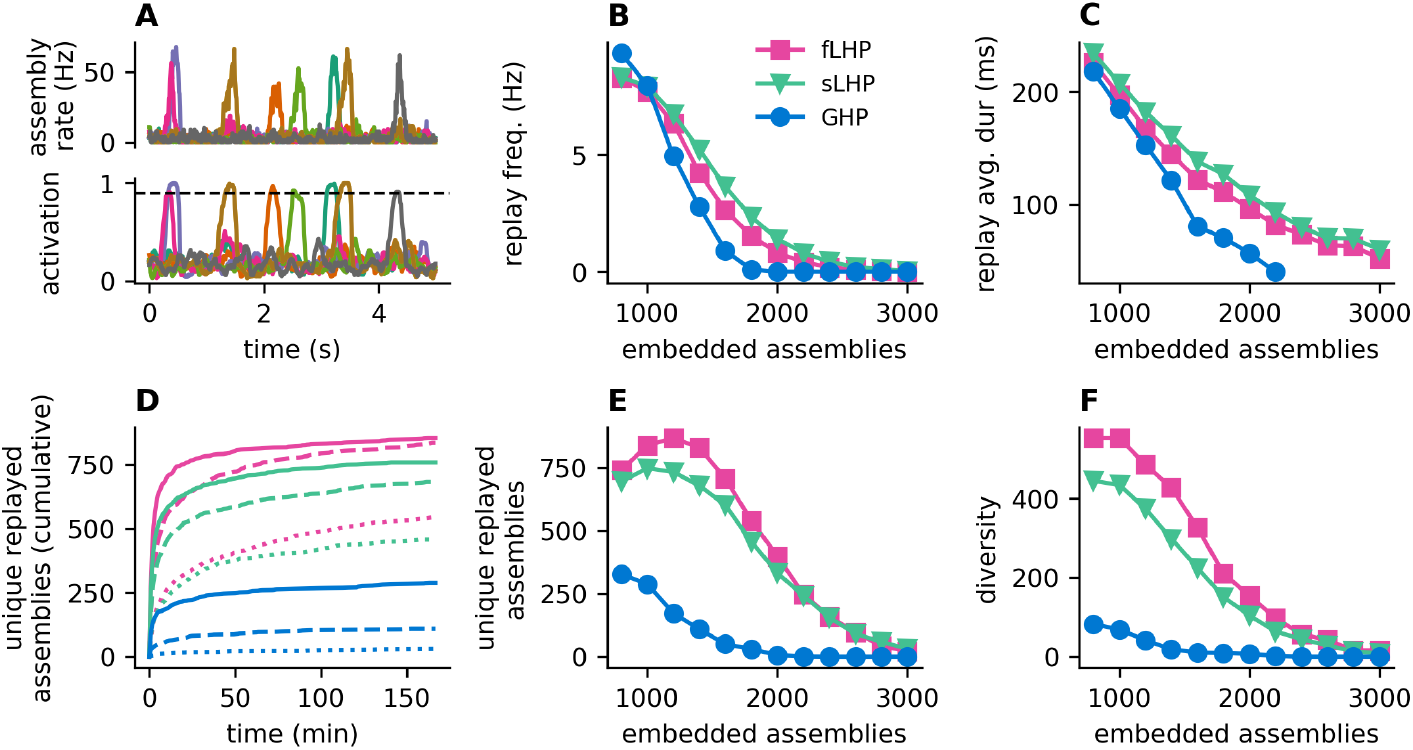
Crucial effects of inhibitory structure on spontaneous replay. **A:** Embedded memory assemblies are spontaneously replayed without informative external input. Top: each colored line represents a firing rate averaged across neurons in a selected assembly. Bottom: the activation of the same assemblies was measured as the fraction of the assembly neurons active within a moving time window. The dashed line represents the threshold beyond which an assembly was regarded as replayed. **B:** The frequency of replay events (the number of events per unit time) decreases with the number of embedded assemblies. **C:** The average duration of a replay event decreases with the number of embedded assemblies. **D:** The cumulative number of unique replayed embedded assemblies during spontaneous activity is plotted as a function of time. Identical assemblies were only counted once. **E:** The number of unique replayed embedded assemblies during 10 000 s of spontaneous activity is plotted. **F:** The diversity of replayed embedded assemblies was estimated with an entropy measure corrected for sampling bias.

We may regard the replay diversity of transient memory states as a measure of storage capacity. The hippocampus likely needs to replay individual assemblies frequently enough during spontaneous activity to consolidate the corresponding memory traces reliably into cortical long-term storage. We evaluated the different inhibitory plasticity rules from the viewpoint of memory capacity. Intriguingly, both the number of unique embedded assemblies that are replayed and the overall diversity of replayed embedded assemblies reveal that networks stabilized with the fLHP rule have the most diverse repertoire of replay activity (Fig. 2D-F). Therefore, the fLHP network can reliably store the highest number of unique memories, yielding the largest memory capacity.

An external stimulus can retrieve an assembly more easily when spontaneous activity replays it more often. Thus, the “replayability” of assemblies is strongly tied to their “retrievability”. To show this, we applied a 100 ms-long external input to a small subset (10) of neurons belonging to an assembly, and measured the proportion of the non-stimulated assembly neurons that fired a spike during the stimulus. We also measured the firing rate of these neurons. The maximum activation and peak firing rate of non-stimulated neurons were strongly correlated with whether an assembly was replayed or not, exhibiting highly skewed or bimodal distributions over memory assemblies (Fig. 3A-I). These results demonstrate a functional link between the replayability and retrievability of memory assemblies.

**Fig. 3.**
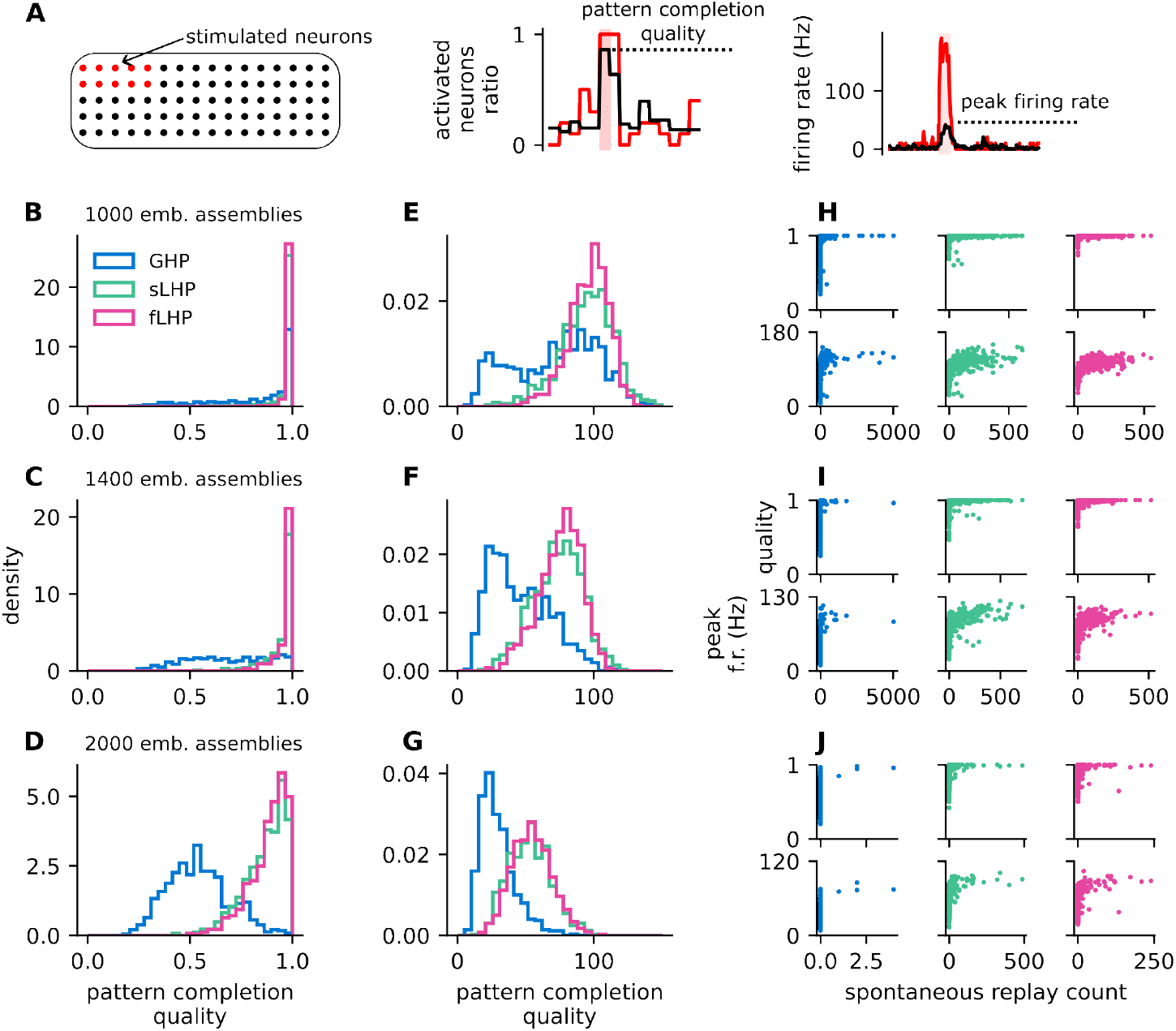
Assembly retrievability is correlated with its replay frequency. **A:** We supplied external stimulus to 10 assembly neurons and measured the response of the remaining, non-stimulated neurons. **B-D:** Histograms of pattern completion quality of non-stimulated assembly neurons during the external stimulation. **E-G:** Histograms of peak firing rate of non-stimulated assembly neurons during the. **G-I:** Dependence of the assembly retrievability on the number of reactivations during spontaneous activity (top rows: pattern completion quality, bottom rows: peak firing rate).

Note that without the presence of SFA the activity of the assembly persisted even after the removal of the stimulus. However, such attractor was typically very unstable and within short time the neural activity switched to a different attractor, which was often a mixture of several different memory assemblies (Fig. S1). This indicates that the network is already operating beyond its conventional capacity for stable memory states.

### Assembly-nonspecific inhibition poorly supports memory replay

The organization of excitatory and inhibitory synaptic inputs (*g*_exc_, *g*_inh_) has a crucial effect on the reactivation of cell assemblies. During spontaneous activity, synaptic inputs to each neuron fluctuate around their central values, with occasional excursions to a high-conductance state, where excitatory conductance is greatly increased in some cell assemblies (Fig. 4A). During such an excursion, neurons in these assemblies are expected to be elevated to an active state from a baseline state. In other words, an assembly is regarded as active when the majority of its member neurons are in the active state. We aimed to clarify what statistical features of the base state significantly contribute to the spontaneous reactivation of assemblies. We found that the multi-state distribution of excitatory and inhibitory conductances biased the statistical estimation of synaptic inputs in the active state (Fig. 4A, purple line). Therefore, we employed a robust covariance estimator (the minimum covariance determinant method, Fig. 4A, red line) for unbiased estimation.

**Fig. 4.**
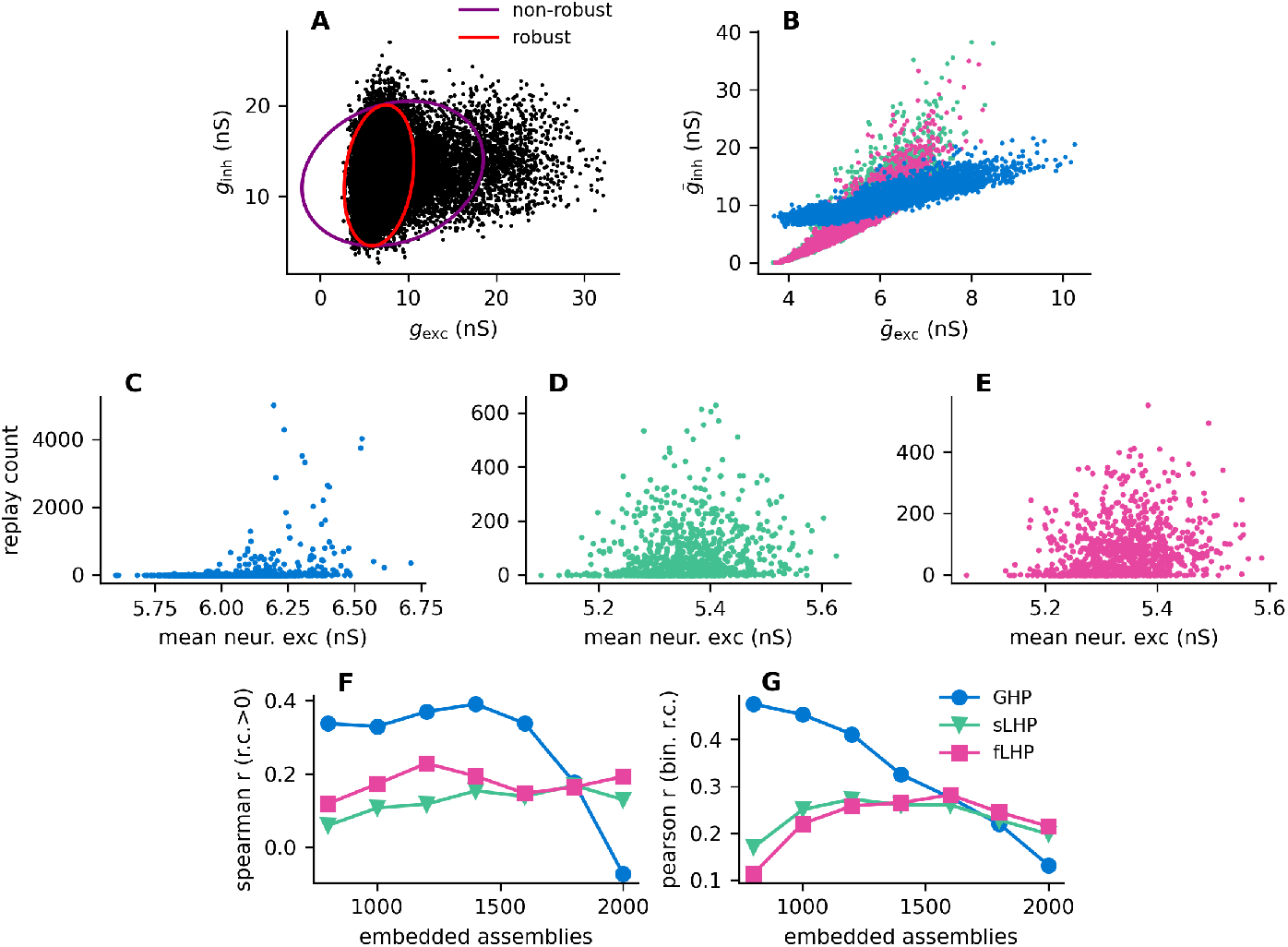
The LHP reduces winner-take-all. **A:** Scatter plot of excitatory and inhibitory conductances during spontaneous activity of an example neuron. The purple ellipse represents moment-based estimation of the covariance matrix, red ellipse the robustly estimated covariance. The ellipse represents the 95th percentile of the estimated distribution. **B:** Robustly estimated mean excitatory and inhibitory input to the excitatory neurons (network with 1000 embedded assemblies). **C:** Spearman’s correlation coefficient of the mean excitatory input and assembly replay count, for assemblies that were replayed at least once. **C-E** Relationship between the mean excitatory input an assembly neuron receives averaged across neurons in each assembly, and how many times the assembly was replayed during a 10 000 s simulation (D: GHP, E: sLHP, F: fLHP).

As a result of the high variability of synaptic weights and the variability in the number of assemblies each neuron participates in, the total excitatory input to an excitatory neuron was highly variable. In all networks the strength of excitatory input and that of inhibitory input (averaged over time) were correlated (Pearson’s r, GHP: 0.89, sLHP: 0.92, fLHP: 0.93, *p <* 10^−5^); neurons receiving strong excitatory input receive strong inhibitory input, and vice versa (Fig. 4B). However, in networks stabilized by the GHP rule, inhibitory inputs are more homogeneous across neurons than excitatory inputs. Consequently, neurons receiving weak excitatory input receive disproportionately strong inhibitory input, and vice versa.

The inhomogeneous excitatory inputs strongly biased the repertoire of replayed embedded assemblies. In all three types of networks, we found positive correlations between the mean excitatory input to an assembly (averaged over time and across assembly neurons) and the assembly replay frequency (Fig. 4C). We evaluated the Spearman’s rank correlation coefficient between the mean excitatory input and replay count for assemblies that were replayed at least once, because a large number of assemblies that were not replayed could bias the rank statistics. The correlation was the highest for the GHP network, up to 1800 embedded assemblies, suggesting that tighter balancing of excitatory and inhibitory inputs helped remove part of the stimulus inhomogeneity in the LHP networks. In the GHP network, assemblies receiving strong excitatory inputs could overcome the inhibitory inputs more easily and became activated more frequently than other assemblies. This implies that cell assemblies compete for spontaneous activation through inhibitory feedback in a winner-take-all (WTA)-like fashion. Many assemblies become suppressed, while a subset of assemblies is activated frequently.

We note that the WTA-like competition does not fully explain the difference in replay diversity between the global (GHP) and local (sLHP and fLHP) rules. With an increase in the number of embedded assemblies, spontaneous replay eventually disappears without yielding any winner. Compared to the locally stabilized networks, replay events disappear at a lower memory load in the GHP network, indicating that the transient attractor structure is substantially weaker in this network.

### Assembly-specific inhibition enables diverse memory replay

The LHP rules organized neural networks to replay more diverse memory assemblies than the GHP rule. Furthermore, the fLHP network replayed a larger number of different memory assemblies compared to the sLHP network. We compared the networks by analyzing their inhibitory structure, specifically whether inhibitory neurons provide input indiscriminately to the excitatory neurons onto which they connect, or are more specialized and provide input to a smaller group of neurons. Similarly to the replay diversity, we used an entropy-based connectivity diversity measure. This diversity measure takes into account the varying outgoing synaptic connections from an inhibitory neuron. It can be interpreted as the effective number of excitatory neurons (Fig. 5A), or the effective number of assemblies (Fig. 5B), the inhibitory neurons connect onto. For instance, the diversity of I-to-E connections of *n* is equivalent to the inhibitory neuron connection to *n* excitatory neurons with the same synaptic weights and synaptic weight zero to all other neurons. Inhibitory neurons in the fLHP network formed I-to-E connections with lower diversity than in the sLHP and GHP networks (Fig. 5A), implying that inhibitory neurons in the fLHP network exhibited a higher degree of specialization. We observed a similar effect when calculating the diversity of the total connection strengths onto the embedded assemblies (Fig. 5B). We also found small but significant differences between the diversity of connections to the embedded assemblies and to random assemblies (shuffle control) (Fig. S2, Tab. S1) in all three networks. This difference was by far the most significant for the fLHP network, indicating that motifs of excitatory and inhibitory neurons significantly contribute to increasing the specialization of inhibitory neurons in this network. This effect, while also present in the GHP and sLHP networks, is negligible compared to that in the fLHP network.

**Fig. 5.**
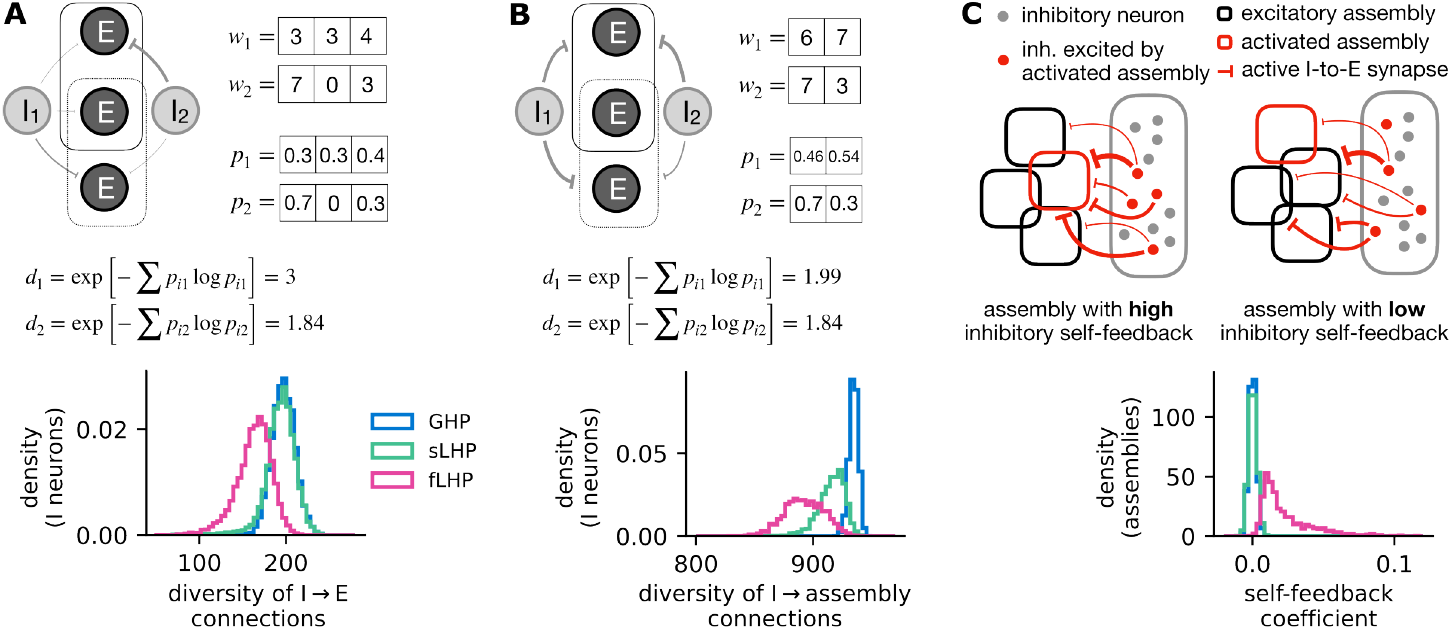
Specialization of inhibitory connections and self-feedback. **A:** We evaluated the diversity of outgoing connections for each inhibitory neuron. Lower diversity means higher specialization. The diagram illustrates the calculation for two inhibitory neurons as an example. **B:** Similar as **A**, but we summed the outgoing synaptic weights for each assembly and calculated the diversity of connections to the embedded assemblies. For comparison with shuffled (non-embedded) assemblies, see Fig. S2. **C:** For each embedded assembly, we calculated how much it inhibits itself by activating the inhibitory neurons, compared to other embedded assemblies.

Furthermore, we specifically searched for E-to-I-to-E loops in the connectivity matrix. For each assembly, we evaluated a self-feedback coefficient, measuring how much the inhibitory feedback inhibits an active assembly compared to non-active assemblies (Fig. 5C; see Methods for details). We found that in the fLHP network assemblies mostly inhibit themselves, while in the sLHP and GHP networks the inhibitory feedback is assembly non-specific.

Finally, we analyzed excitatory and inhibitory conductances on excitatory neurons during spontaneous replay in all three cases of inhibitory structure to see how this structure translates into an effect on the neural dynamics. We averaged the traces of excitatory and inhibitory conductances for embedded assemblies over their replays during 100 s of spontaneous activity. The average was taken across assembly neurons and all replayed embedded assemblies. As expected, excitatory input to assembly neurons grew several-fold during an assembly replay (Fig. 6A). Inhibitory input to a replayed assembly also increased in all three networks. However, we found that in the GHP and sLHP networks, inhibitory inputs to currently non-replayed assemblies grew in parallel with those to currently replayed assemblies. In contrast, in the fLHP network, inhibitory input to a replayed assembly was stronger than that to the remaining assemblies (Fig. 6B-C). Thus, the specialization of inhibitory neurons arising from the connectivity matrix dramatically modifies network dynamics during assembly replay.

**Fig. 6.**
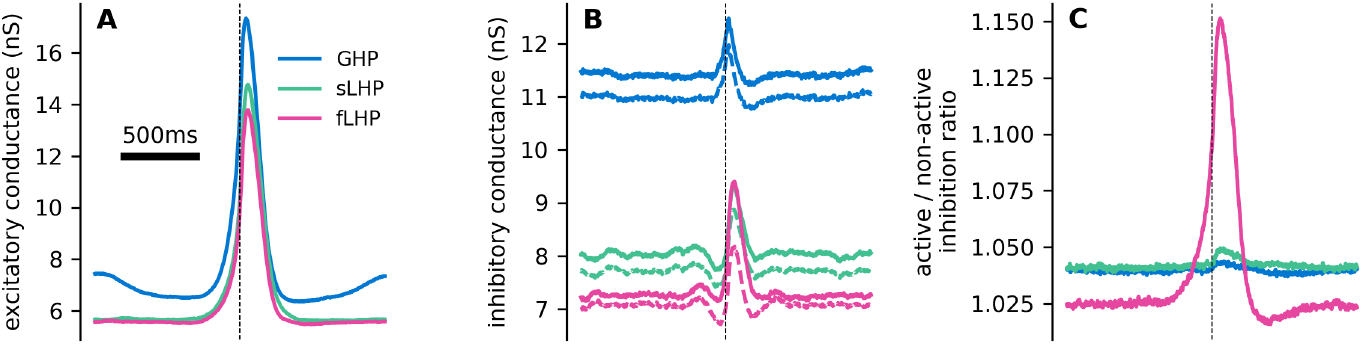
Self-feedback during replay. **A:** Excitatory conductance of the replayed embedded assembly (averaged across the assembly neurons) for the three different rules. **B:** Inhibitory conductance of the replayed embedded assembly (averaged across the assembly neurons) for the three different rules (full lines) and the average inhibitory conductance of other assemblies during the replay (dashed lines). In each case, both inhibitory input to the replayed embedded assembly and to the other assemblies increases during the replay. Note that the oscillations before and after the peak are an averaging artifact arising from lower probability of two different activations being close to each other. **C:** Ratio of the inhibitory input to the replayed embedded assemblies and the input to the remaining assemblies. In the fLHP network, input to the replayed embedded assembly is about 10% larger than to the other assemblies, indicating assembly-specific inhibition. Traces were obtained by averaging 1552, 1557, and 1495 spontaneous replays in the GHP, sLHP, and fLHP networks respectively, with 1000 embedded assemblies. Dashed line indicated the time of detected replay start, at which individual traces were aligned.

### Effective dynamical modes of excitatory neurons

We now turn to exploring the network mechanisms of self-organizing the differential memory replay in the three network models. Although synaptic plasticity only modified the I-to-E connections but not the other connection types, the changes in the overall network dynamics could change the sensitivity (susceptibility) of individual neurons to synaptic input. We linearized the network dynamics around the baseline activity and derived an effective connectivity matrix, which takes the sensitivity of each neuron into account (Fig. 7A). The linearized network obeys the following dynamics:

**Fig. 7.**
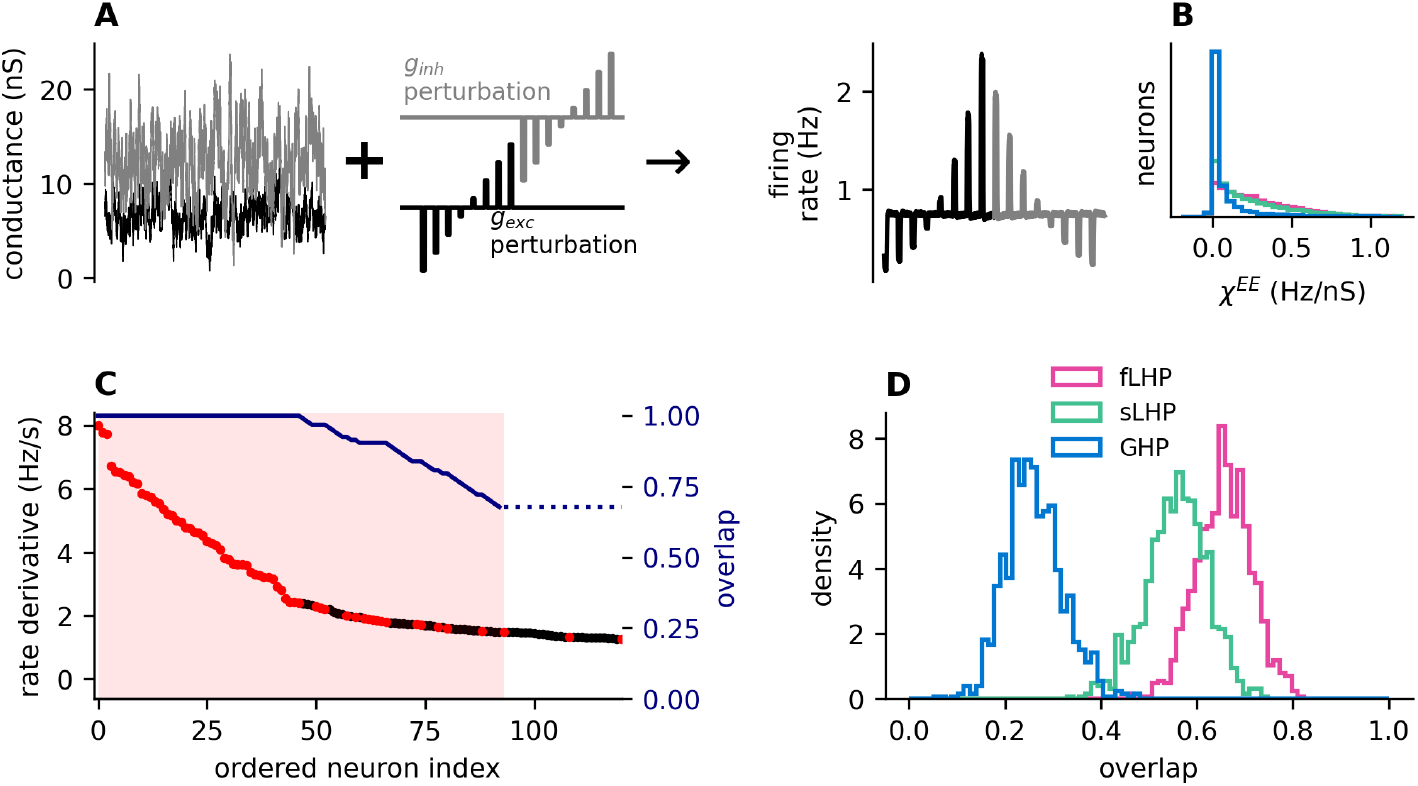
Alignment of embedded assemblies in a linear model. **A:** Estimation of neuron’s sensitivity to synaptic input. For each neuron we replace the input from the network with two correlated Orstein-Uhlenbeck processes with means and covariance matching the robust estimator (black represents excitatory input, gray inhibitory input). On top of the background input we added small perturbation in the excitatory or inhibitory conductance and measured the firing response of the neuron to these perturbations. **B:** Distribution of susceptibilities of excitatory neurons to excitatory synaptic input. **C:** We estimated how the *EE* portion of the effective connectivity matrix maintains the direction of the assembly vectors ***ξ***_***µ***_. We multiplied 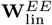 and calculated the overlap between the most affected neurons and the assembly neurons. The shaded portion represents the size of the assembly *µ*. Dots are the largest 120 values of 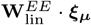 in descending order. Red dot means the neuron belongs in the assembly ***ξ***_***µ***_, black otherwise. The black line indicates what fraction of assembly neurons is present within the top *n* neurons of the derivative. Line ends where *n* is the size of the assembly *s*(*µ*), where the final overlap value is evaluated. **D:** Histograms of the overlap values in networks with 1000 embedded assemblies. GHP stabilized network shows significantly smaller overlap than sLHP and fLHP stabilized networks.

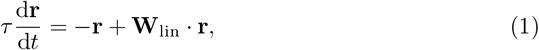

where **W**_lin_ is the effective connectivity matrix, i.e., the Jacobian at the base state, in which no neurons are in an activated state, obtained by multiplying the synaptic weight matrix by the sensitivity of the post-synaptic neurons to synaptic input (see Eqs. 32–33 in Methods).

Overall, neurons in sLHP and fLHP networks were much more sensitive to synaptic input than neurons in the GHP network (Fig. 7B, Fig. S3A), in which the variability of sensitivity across neurons (measured as the coefficient of variation, Fig. S3B) was also higher. The lower sensitivity in the GHP network may be most conveniently explained by the lower resting potentials of neurons (Fig. S3). We also note that, in the sLHP and fLHP networks, the resting potentials decreased with the mean excitatory input due to a strong compensating inhibitory input. In contrast, in the GHP network, the resting potentials and, subsequently, the sensitivity increased with the mean excitatory input.

We studied how the I-to-E plasticity affects assembly embedding in the E-to-E connectivity. To this end, we analyzed how well the submatrix 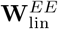 preserves memory traces, i.e., assembly vectors 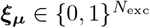, where 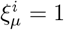 or 0 if excitatory neuron *i* belongs or does not belong to the assembly *µ*, respectively. Then, the vector 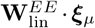 represents the derivative of the linearized system retrieving the memory assembly *µ*. To study the robustness of the transient memory states, we looked at the overlap of the *s*(*µ*) most active neurons of 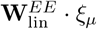 with non-assembly neurons (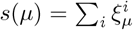 is the size of the assembly *µ*; overlap of 1 means the most active neurons are all neurons from the assembly *µ*; Fig. 7C). We found that the overlap was substantially higher for the sLHP and fLHP rules compared to the GHP rule (Fig. 7D). The results demonstrate that the LHP rules modify I-to-E connections such that the effective E-to-E connections can embed the assemblies more reliably. Further, we tested the ability of pattern completion by calculating the overlap of 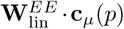 with *ξ*_*µ*_, where **c**_*µ*_(*p*) is a corrupted version of the *µ*-th memory pattern in which that 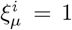 for neurons in the assembly with probability *p*, 0 otherwise (*c*(*ξ*_*µ*_; 1) = *ξ*_*µ*_) and found that the fLHP linearized network performs better than the sLHP linear network, which in turns performs better than the GHP linearized network (Fig. S4).

The GHP network is stabilized in a different manner than sLHP and fLHP networks are. To analyze the stability of these networks, we calculated the eigenspectra of the effective connectivity matrices (Fig. 8A). With 1000 embedded memory assemblies, all three networks had eigenvalues mostly contained within a circle of radius of approximately 0.5 (with the exception of a large negative real eigenvalue around -5). However, with increasing memory load, the eigenspectrum of the GHP network became elongated. We looked at the eigenvector corresponding to the largest eigenvalue and selected the top 50 neurons based on this eigenvector (Fig. 8B). We considered the set of these neurons as a new, eigenvector-based assembly. This eigenvector-based assembly was a mixture of embedded assemblies, but the largest overlap with any of the embedded assemblies was only 10 neurons (Fig. 8C). We observed that the eigenvector-based (non-embedded) assembly was periodically spontaneously activating, but the embedded assemblies that had the largest overlaps with the eigenvector-based assembly did not fully activate (Fig. 8D). This indicates that the eigenvector is dominating the dynamics of the network and preventing the embedded assemblies from activation, presumably through lateral inhibition. In contrast, we saw no such effects with an eigenvector-based assembly identified from the largest eigenvalue in the fLHP network (Fig. 8E-G). This implies that the fLHP network activates embedded assemblies with much smaller mutual interference.

**Fig. 8.**
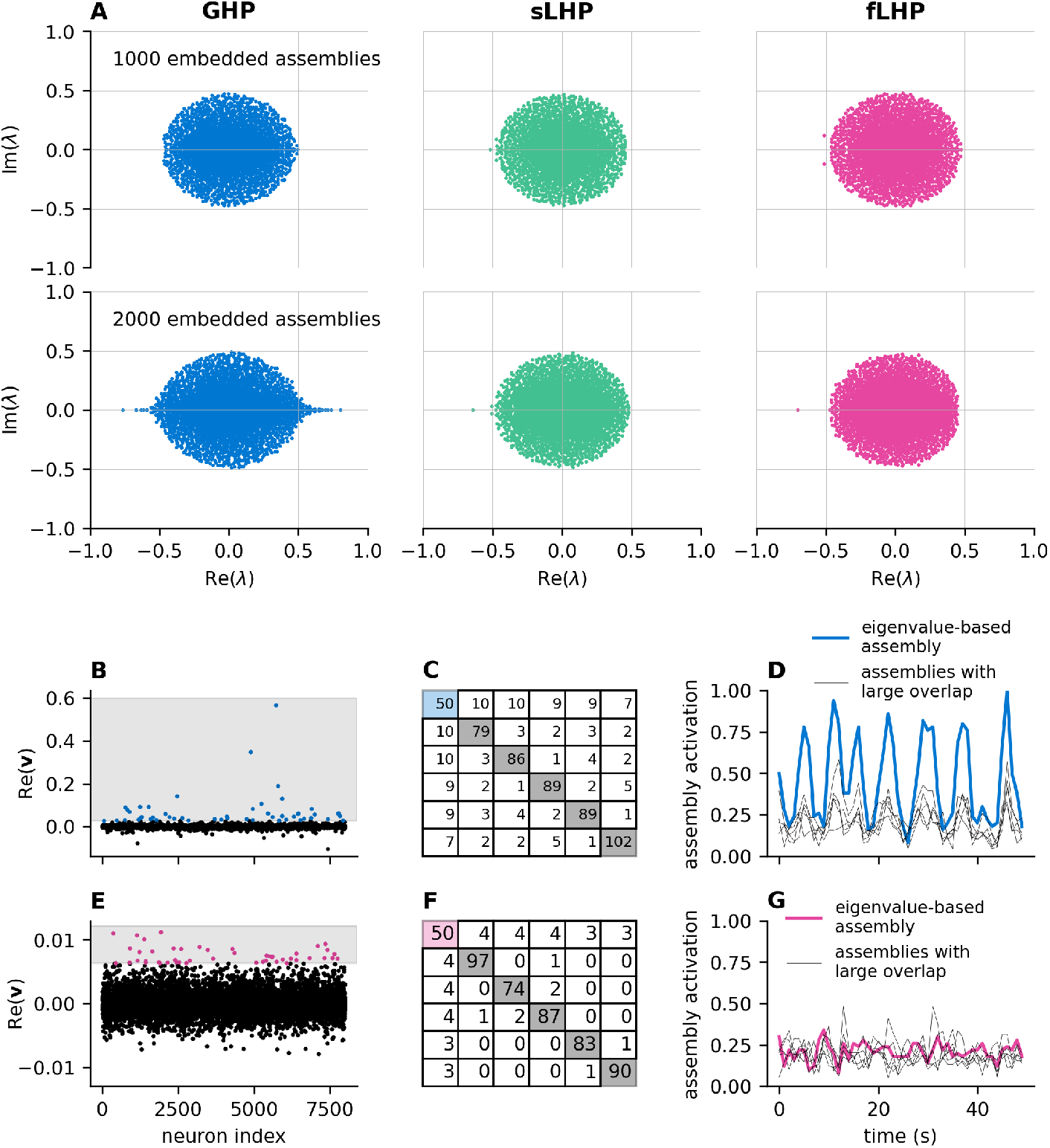
Eigenspectra of linearized models. **A:** Eigenspectra of Jacobians of linearized models. Note that in each case, the spectrum also contained a large negative value around -80, which is not displayed for compactness. The shape of the eigenspectra does not change greatly with the number of embedded assemblies for the sLHP and fLHP stabilized model, but the spectrum of the GHP stabilized model begins to elongate and its spectral radius increases. **B:** Excitatory part of the eigenvector corresponding to the eigenvalue with the largest real part of the GHP model with 2000 embedded assemblies. Shaded area shows the threshold used to identify a new, eigenvalue-based assembly, consisting of 50 neurons. Color identifies which neurons exceed that threshold. **C:** Table showing the overlap of the eigenvalue-assembly with the 5 assemblies with largest overlap. Sizes of the assemblies are on the diagonal, non-diagonal terms are the numbers of shared neurons between the two assemblies. This table illustrates that the eigenvalue assembly is a mixture of several different assemblies, but without any one of them dominating the others. **D:** Activation of the eigenvalue-based assembly (in blue). Gray lines represent 5 embedded assemblies that have the largest overlap with the new eigenvalue-based assembly. Activation is the fraction of neurons in an assembly that fire a spike in a 100 ms interval. **E-G:** Same as (B-D), but for the fLHP network with 2000 embedded assemblies.

## Discussion

Active roles have been suggested for transient memory states experimentally and theoretically [35, 36], but the mechanism by which hippocampal networks replay memory traces spontaneously and transiently remains elusive. As activity replay is crucial for memory consolidation, which is the primary role of the hippocampus, we constructed a spiking neural network that is capable of spontaneously replaying embedded memory assemblies. Due to spike-frequency adaptation, memory assemblies exhibited only transient, but not persistent, activity in the model. We examined how such transient activity enables the neural network to replay more assemblies and enhance its potential capacity for memory consolidation. To this end, we tuned the stability of spontaneous network activity by using global and local inhibitory plasticity rules. These rules stabilized network activity differently, where the local rules produced an assembly-specific E-I balance advantageous for diverse memory replay.

### Comparison between different inhibitory plasticity rules

In spiking neural networks, the stabilization of network activity is crucial and is believed to rely significantly on plasticity at inhibitory synapses. Our results suggest that the biological details of inhibitory stabilization heavily affect the efficiency of memory storage in a recurrent network. The GHP rule induces a global network-level homeostasis. Neurons receiving stronger excitatory inputs formed stronger assemblies that were replayed more often than others, implying that the replay opportunity was biased toward such assemblies. In contrast, neuron-specific homeostasis induced by the LHP rules significantly enhanced replay diversity by balancing neurons receiving stronger excitation with stronger inhibitory feedback. This prevents such neurons from dominating the network, allowing a uniform replay of various assemblies.

Assuming the different timescales of local homeostatic plasticity, we could obtain two types of inhibitory structure. The fLHP rule [27] is biologically plausible and has been experimentally observed in different brain areas [22, 23]. This rule achieves cotuning of excitation and inhibition, which is consistent with experimental observations [37–39]. For comparison, we introduced the sLHP rule to achieve local homeostasis without a tight co-tuning of excitation and inhibition. This rule is identical to the fLHP rule except for a significantly longer time constant. In general, both local homeostatic rules led to far more diverse replay than the GHP rule. However, the network tuned on the faster timescale was able to replay with higher diversity than the network tuned on the slower timescale. This result highlights the crucial role of finely timed co-tuning of neuron-specific E-I balance in organizing diverse replay.

While our models exhibited no evidence of catastrophic forgetting, the overall frequency of memory replay decreased with an increase in the number of embedded memory assemblies, signaling overall weakening of the attractor structure. The replay frequency dropped much more slowly in the LHP networks than in the GHP network.

This indicates that the effect of LHP rules is twofold: these rules allow more assemblies to be replayed more often. We explained the attractor structure in the individual networks through linear rate approximations of spiking networks and the resultant effective E-to-E connection matrix. The leading eigenvectors of this matrix enabled us to identify assemblies of spontaneously active neurons. We found that, despite synaptic weights between excitatory neurons being fixed, the LHP rules modify the effective E-to-E connectivity to change their sensitivity to synaptic input in such a way that the E-to-E submatrix embeds memory assemblies more strongly.

The effective matrix also showed interesting differences between the inhibitory plasticity rules. In the GHP network, increasing the number of embedded assemblies makes the network dynamics dominated by the leading eigenvectors, which were generally a mixture of the embedded memory assemblies. These assemblies are reminiscent of the so-called spin-glass states that occur in Hopfield-type auto-associative memory networks and dominate their retrieval dynamics when the memory load exceeds the critical capacity [40]. We speculate that the activation of these eigenvector-based mixture-assemblies prevents the embedded assemblies from spontaneously activating. In contrast, we could not identify any eigenvector-based assembly in the sLHP and fLHP networks, at least for the range of memory load examined in this study.

### Implications of spontaneous replay for catastrophic forgetting

For fixed lognormal E-to-E synaptic connections, assembly replay becomes less frequent and less diverse with more assemblies embedded in E-to-E synapses. Intriguingly, this decrease in replay diversity is gradual and not accompanied by the abrupt disappearance of transient memory states [41–45]. This implies that, unlike in stable attractor-based models, forgetting may not be a “catastrophe” in neural networks replaying transient memory states.

Indeed, transient memory states mitigated catastrophic forgetting. In our model, removing the spike-frequency adaptation turned transient activity into sustained activity in some assemblies. However, this sustained activity was typically short-lived and quickly transitioned into a state sustaining another memory assembly or a mixture of different memory assemblies. It is said that a network with transient memory states can reinstate the embedded memory states beyond the memory load at which the corresponding network with persistent memory states ceases to function properly.

### Limitations of the models and open questions

In our work, we employed fixed E-to-E connections without considering any form of excitatory plasticity. Similar to previous studies exploring the limits of memory storage [9, 46], we assumed that memory assemblies were hard-wired beforehand. Fixed E- to-E connections allowed us to embed an arbitrary number of memory assemblies in the network. However, regardless of inhibitory plasticity rules, we speculate that a network exhibiting spontaneous replay generally has a critical memory load at which replay diversity is maximized. This point needs further investigation.

Other studies have investigated the formation of assemblies in networks subject to excitatory plasticity and the GHP and fLHP inhibitory plasticity rules [28, 30, 47]. Hybrid approaches have also been proposed for combining different plasticity rules between subnetworks to stabilize network activity [48, 49]. Clarifying whether networks with higher replay diversity are capable of maintaining more memories is an intriguing open question. Combining I-to-E plasticity with E-to-E plasticity, we may further elucidate the mechanism of co-tuning excitatory and inhibitory assemblies for spontaneous replay.

Theoretical studies suggest that inhibitory engrams that may arise from fine co-tuning of excitation and inhibition [50] offer certain advantages, such as recall of memories through disinhibition [31]. Inhibitory assemblies have also been shown to promote a winnerless competition in a network of binary neurons [51], which aligns well with our results. Our result that assembly-specific (fLHP) inhibition improves replay diversity is also in agreement with the previous finding that the dominance of assembly-specific inhibition over assembly-non-specific inhibition suppresses the WTA-like behavior [52]. It would be interesting to investigate whether increasing the stimulus-specificity of inhibitory cells would further increase replay diversity.

Finally and most importantly, the present model contains only a single type of inhibitory neuron. However, in reality, multiple inhibitory cell types exist in cortical circuits to serve different computational demands. For example, a computational study has shown that inhibitory neurons acting on axon terminals, which is a characteristic of parvalbumin (PV) neurons, can help with rapid stabilization during learning [53]. Another study has demonstrated computationally that somatostatin neurons exhibiting asymmetric Hebbian plasticity may increase competition between assemblies [23]. For the sake of simplicity, we focused on inhibitory neurons with symmetric Hebbian plasticity, which could be assigned to PV neurons [23]. Exploring how the different inhibitory neuron subtypes orchestrate to maximize the replay performance is especially important.

## Methods

### Neuron model

We modeled leaky integrate-and-fire neurons with conductance-based synapses

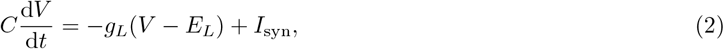

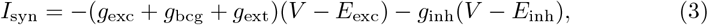

where *C, V, g*_*L*_, and *E*_*L*_ are the membrane capacitance, membrane potential, leak conductance, and leak reversal potential, *g*_exc_ and *g*_inh_ the synaptic conductances and *E*_exc_ and *E*_inh_ the respective reversal potentials, *g*_bcg_ is the conductance from a background input the CA3 receives from the entorhinal cortex, and *g*_ext_ is the external input which is supplied during the cued retrieval experiment (Fig. 3). Neurons fire a spike when the membrane potential reaches a dynamic threshold *θ*(*t*), modeled as in the MAT model [54]:

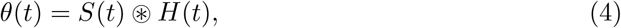

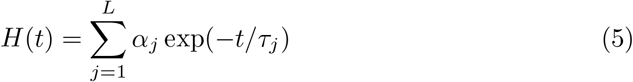

where *k* iterates over all previous spikes fired by the neuron. This dynamic threshold provides SFA on *L* different time-scales *t*_*j*_. In contrast to the MAT model, upon firing a spike the membrane potential *V* is reset to the value *E*_*L*_.

Synapses are exponential:

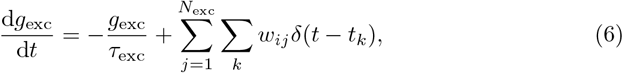

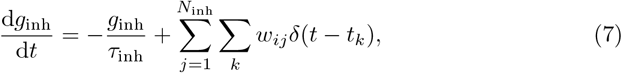

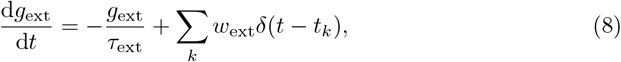

with *w*_*ij*_ being the synaptic strength of connection from neuron *j* to neuron *i, k* iterates over all spikes of the *j*-th neuron or spikes supplied during external stimulation for cued retrieval. During the cued retrieval, each stimulated neuron received Poisson input from an independent external population with rate *λ*_ext_ = 2 kHz and synaptic weight *w*_ext_ = 0.3 nS for 100 ms.

Due to the lognormal distribution of the synapses, as described below, the network was typically capable of sustaining spontaneous activity without any external input[32, 55]. However, the spontaneous activity typically terminated during longer simulations. In order to ensure that the network remains active for long periods The background input is modeled independently for each neuron as:

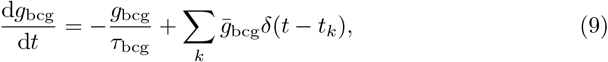

where *t*_*k*_ are spike times generated from an independent Poisson process with frequency *λ*_bcg_ and 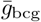 is the conductance amplitude. The neuron model parameters are provided in the Tab. 1.

**Table 1.**
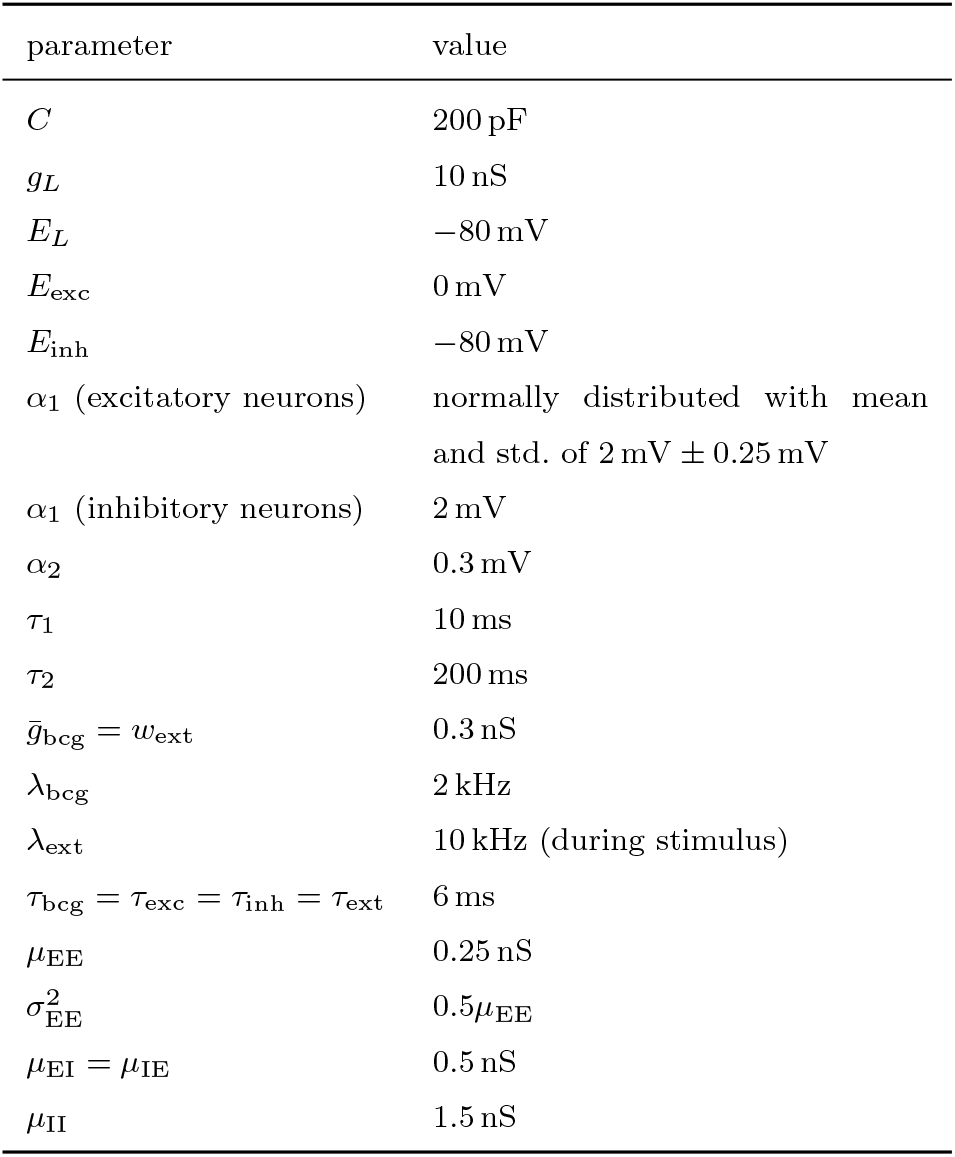
Parameters of the neuron model and synaptic weights.

### Network model

We modeled a network of 8000 excitatory (E) and 2000 inhibitory (I) neurons. We generated random assemblies, each excitatory neuron belonged to a pattern with probability *p*. We used *p* = 0.01, making the assemblies consist of ensembles of 80 neurons on average.

To set the E to E connections, we used an approach similar to Mongillo et al. [46] and Hiratani and Fukai [9], with the main difference being that only excitatory neurons were a part of a memory pattern, and we did not exclude symmetric connections due to their abundance in the CA3 [56]. In particular, we first calculated the Hebbian terms of the E to E connections as:

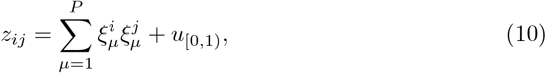

where 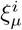 is 1 if neuron *i* belongs to the memory pattern *ξ*_*µ*_ and 0 otherwise, *P* the number of memory assemblies, *u*_[0,1)_ is a random number from the uniform distribution between 0 and 1. Next, we only kept the connections within top 5% of the Hebbian terms. To these connections we assigned synaptic weights 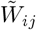 drawn from a lognormal distribution with mean *µ*_EE_ and variance 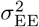 in the same order as the Hebbian terms:

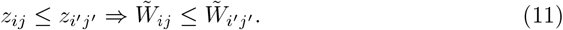

Next, to prevent the appearance of excitatory hubs destabilizing the network, we followed Hiratani et al. and normalized synaptic weights over pre-synaptic neurons as:

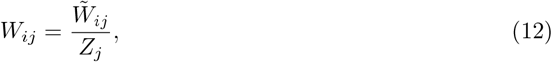

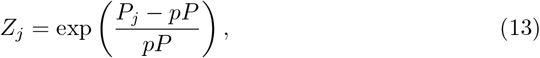

where *P*_*j*_ is the number of assemblies that the pre-synaptic neuron *j* participates in, and *pP* the expected number of assemblies that any neurons participates in.

All connections other than E to E connections were randomly selected to be nonzero with 10% probability and assigned a constant value *µ*_XY_ where X is the post-synaptic population and Y is the pre-synaptic population. See Tab. 1 for values of synaptic weights. Each synaptic connection is assigned a delay between 1 ms and 3 ms sampled from a uniform distribution. The above-presented method assumes that the excitatory assemblies have both higher probability of connection within an assembly and stronger synaptic connections (Tab. 2).

**Table 2.** Probability of connection and average synaptic strength between two excitatory neurons within an assembly, vs. within random neurons.

| | pat. conn. prob. | rand. conn. prob. | pat. avg. $W_{ij}$ (nS) | rand. avg. $W_{ij}$ (nS) |
| --- | --- | --- | --- | --- |
| 800 | 0.66 | 0.05 | 0.31 | 0.25 |
| 1000 | 0.54 | 0.05 | 0.33 | 0.25 |
| 1200 | 0.47 | 0.05 | 0.34 | 0.25 |
| 1400 | 0.42 | 0.05 | 0.35 | 0.25 |
| 1600 | 0.38 | 0.05 | 0.35 | 0.25 |
| 1800 | 0.36 | 0.05 | 0.35 | 0.25 |
| 2000 | 0.34 | 0.05 | 0.35 | 0.25 |
| 2200 | 0.33 | 0.05 | 0.35 | 0.25 |
| 2400 | 0.32 | 0.05 | 0.35 | 0.25 |
| 2600 | 0.31 | 0.05 | 0.34 | 0.25 |
| 2800 | 0.30 | 0.05 | 0.33 | 0.25 |
| 3000 | 0.30 | 0.05 | 0.33 | 0.25 |

### Synaptic plasticity

To achieve a target mean firing rate of the network (GHP) we used the learning rule relying on a global secreted factor *G* [28]:

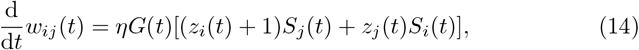

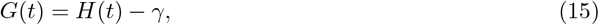

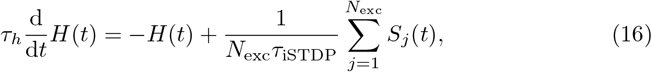

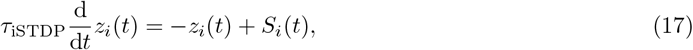

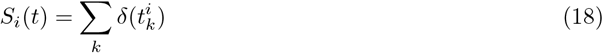

where *γ* is the target firing rate, *G*(*t*) is the global secreted factor tracking if the average firing rate is too high or too low, and 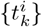 are spiking times of the *i*-th neuron. This learning rule maintains the present heterogeneity in the firing rates of individual neurons, but enforces the firing rate averaged across all excitatory neurons to be *γ*.

In order to achieve microscopic EI balance we used the homeostatic plasticity rule [27]. The I-to-E synaptic weight is modified when either the pre-synaptic or post-synaptic neuron fires a spike:

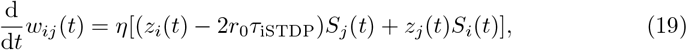

where *r*_0_ is the target firing rate of the post-synaptic neuron. We varied the time constant *τ*_iSTDP_ to switch between neuron-specific (sLHP, *τ*_iSTDP_ = 10 s) and neuron- and-time-specific (fLHP, *τ*_iSTDP_ = 20 ms) rules.

We trained the models for 2000 s with a target firing rate 3 Hz and learning rates 10^−3^ for the GHP and fLHP rules and 2 × 10^−5^ for the sLHP rule. After the training we set *η* = 0.

### Network structure analysis

To evaluate the specialization of inhibitory neurons, we used an entropy-based measure. For each inhibitory neuron, we calculated the entropy of its synaptic weights and the diversity as the exponential of the entropy:

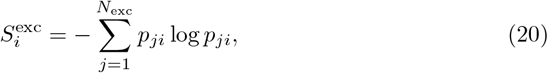

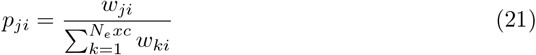

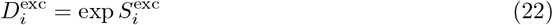

Note that we treat *p*_*ji*_ log *p*_*ji*_ as 0 when *p*_*ji*_ = 0.

Similarly, we calculated the diversity for connections to the excitatory assemblies:

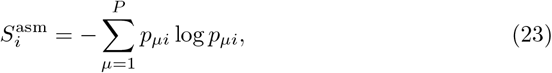

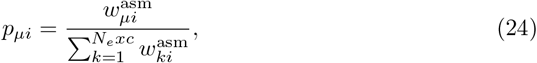

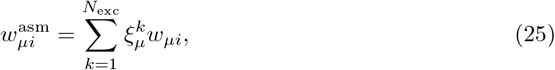

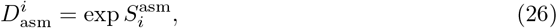

where 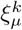 is 1 if the *k*-th excitatory neuron belongs to the assembly *µ* and 0 otherwise.

To calculate the self-feedback coefficient, we first calculated a matrix summarizing the inhibitory effect of assemblies on each other:

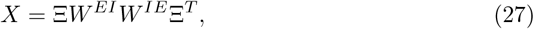

where Ξ is a matrix of assembly vectors, *i*-th row is *ξ*_*i*_. Next, to control for some assemblies receiving more inhibition in general, we obtained *X*^*′*^ by dividing each row of *X* by the row average, but leaving out the diagonal terms:

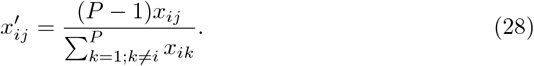

Finally, we calculated the feedback coefficient as the difference between the normalized feedback input of an assembly onto itself 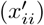 and the averaged normalized inhibitory feedback input to other assemblies 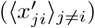.

### Linear rate network approximation

In order to understand the effects of different plasticity rules on the excitability of neurons, we approximated the spiking neural network as a system of linear rate neurons. We performed this in several steps:

1. We extracted the statistics of excitatory and inhibitory input conductances.
2. We simulated each neuron without any presynaptic connections, but with a stochastic input reproducing the statistics of the network.
3. By perturbing the excitatory and inhibitory input and measuring the response of the neurons, we estimated the neurons’ excitability.
4. We combined the neurons’ excitability values with the synaptic strengths to obtain the effective connectivity matrix of the linear system.

To obtain the input statistics, we first sampled the input conductances *g*_exc_ and *g*_inh_ with 1 kHz frequency during 20 s of spontaneous activity. We estimated the mean and variance of *g*_exc_ and 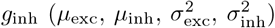 and their correlation coefficient *ρ* with the minimum covariance determinant estimator implemented in Scikit-learn (MinCovDet) to avoid outliers in conductance during pattern activation [57]. We then reproduced these statistics individually for each neuron by replacing *g*_exc_ + *g*_bcg_ and *g*_inh_ with two correlated Ornstein-Uhlenbeck processes:

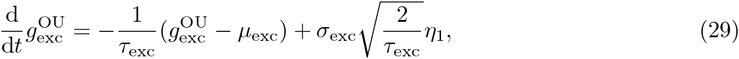

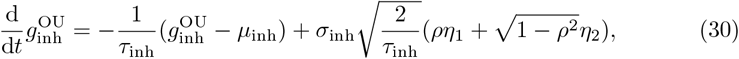

where *η*_1_ and *η*_2_ are two independent realizations of Gaussian white noise. We then modeled the synaptic current in Eq. 2 as:

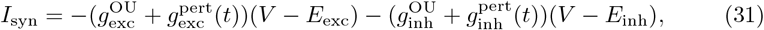

where 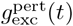 and 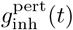 are perturbative inputs used to determine the excitability. The neurons received input of 8 different values (−0.35 nS, −0.25 nS, −0.15 nS, −0.05 nS, 0.05 nS, 0.15 nS, 0.25 nS, 0.35 nS) for 100 ms with 400 ms gaps between two subsequent stimuli. We calculated the mean firing rate during the 100 ms across 1000 trials and fitted a 3rd degree polynomial to the spiking responses of each neuron. We took the first derivative at 0 nS multiplied by 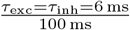 (to account for the short time constant of the synapse) as the excitability of the neuron. Note that an average synaptic weight of 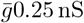 with exponential time constant *τ* 6 ms and firing rate *λ* above base state will result in an average increase in synaptic input of 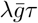. Assuming the values 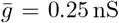 and *τ* = 6 ms, an average increase in conductance by 0.05 nS will be reached with and input of 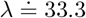 hertz.

We performed the sensitivity estimation separately for excitatory and inhibitory inputs, thus obtaining for each neuron its sensitivity to excitatory and inhibitory input (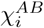, where *i* and *A* mark the index and subpopulation of the post-synaptic neuron, *B* marks if the presynaptic input is excitatory or inhibitory). For each neuron, we then multiplied the presynaptic excitatory and inhibitory weights in the synaptic weight matrix *W* by the neuron’s sensitivity to excitatory and inhibitory synaptic input respectively:

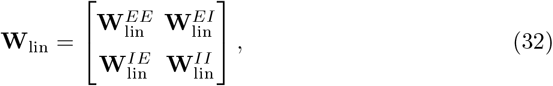

where

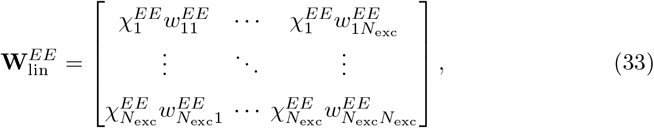

and similarly for other blocks.

### Measuring replay diversity

To evaluate the replay diversity, we used an entropy-based metric. We first converted the replay counts (*r*_1_, *r*_2_, …, *r*_*P*_) into a probability distribution (*p*_1_, *p*_2_, …, *p*_*P*_), where

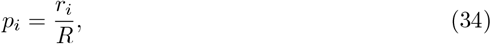

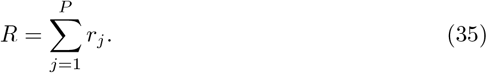

We then calculated the entropy of the distribution *S* and diversity *D* as

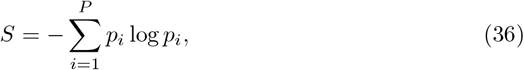

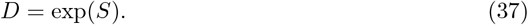

Estimating entropy from a sampled probability distribution introduces a negative bias in the estimated entropy. To obtain an unbiased estimate *Ŝ*, we assumed the following dependency of estimated biased entropy *S*_*k*_ on the total number of observed replays *k* [58]:

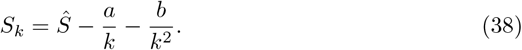

We calculated *S*_*k*_ for different time points of the simulation (2500 s, 5000 s, 7500 s, 10 000 s), obtained *Ŝ* from a non-negative linear regression, and finally obtained the corrected diversity as 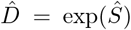, which is presented in the Fig. 2. If no replay happened by the 2500 s mark, we set *Ŝ* = 0. Note that the corrections were typically minimal, implying that 10 000 s of spontaneous activity simulation was sufficient (Tab. S2).

## Data availability

Numerical data necessary to replicate the figures are available at https://github.com/oist-ncbc/bartafukai2025.

## Code availability

All codes for numerical simulations and data analysis tools used in this study are available at https://github.com/oist-ncbc/bartafukai2025.

## Acknowledgements

We thank Kaoru Inokuchi and Kazumasa Tanaka for their valuable comments and suggestions on our manuscript. This study was supported by JSPS KAKENHI no. JP23H05476 to T.F.

## Conflicting interests

The authors declare no competing interests.

## Extended Data

**Fig. S1.**
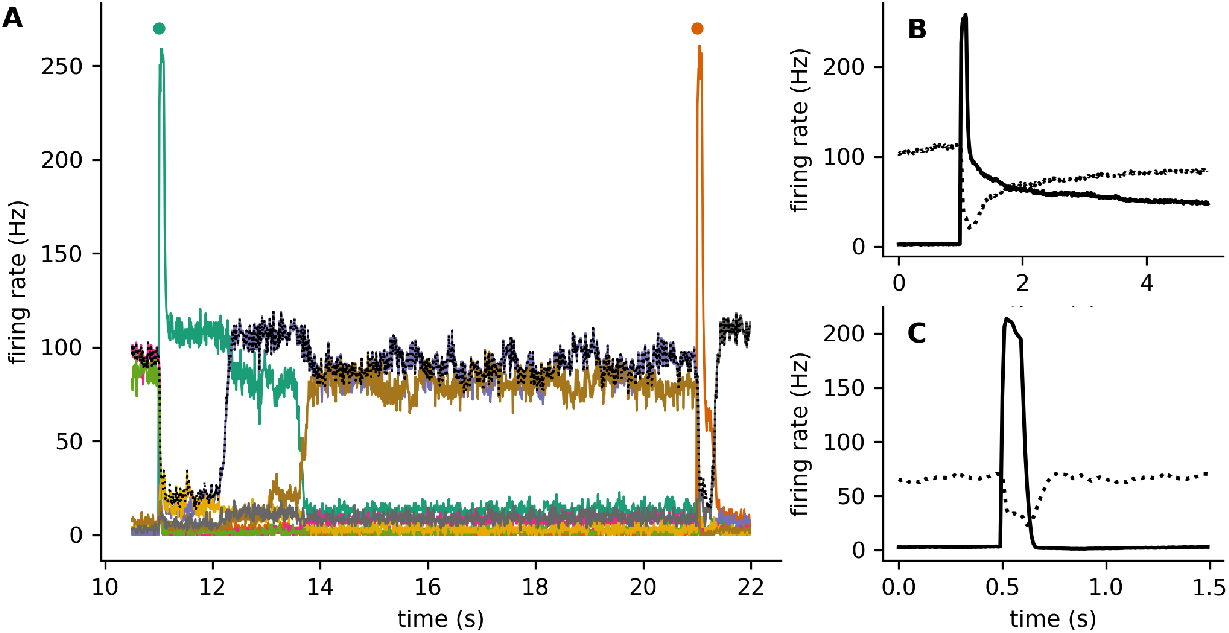
Transient activity allows more reliable recall. **A:** Assembly activity in networks without SFA. A different assembly was stimulated every 10 s, indicated by the colored dot. Different colored lines indicated the firing rate of neurons averaged across different assemblies. Dotted black line marks the maximum firing rate at each time point across all assemblies (with the exception of the two assemblies stimulated in the selected time window). **B:** Firing rate of stimulated assembly in a non-SFA network averaged across stimulated assemblies (full line). Dotted line indicates the assembly with the highest firing rate at each time point, across the remaining assemblies. **C:** Same as **B**, but for the network with SFA.

**Table S1.** Specialization with embedded and random assemblies. We compared the entropy of inhibitory weights across embedded assemblies 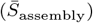 with entropy across random assemblies 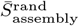. The table shows the relative difference of means 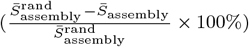. Negative difference means that inhibitory neurons are more specialized on the embedded assemblies, than on random assemblies. This effect is most pronounced with the fLHP rule. *T* statistic from a paired t-test and the corresponding *p*-value are shown. The test had 1999 degrees of freedom in all instances (2000 inhibitory neurons).

| Stabilization rule | embedded assemblies | $\bar{S}_{\text{assembly}}$ | rel. difference (%) | $T$ | $p$-value |
| --- | --- | --- | --- | --- | --- |
| fLHP | 800 | 6.56 | -0.16 | -51.51 | $< 10^{-100}$ |
| | 1000 | 6.79 | -0.11 | -46.43 | $< 10^{-100}$ |
| | 1200 | 6.99 | -0.06 | -30.43 | $< 10^{-100}$ |
| | 1400 | 7.15 | -0.04 | -21.20 | $< 10^{-89}$ |
| | 1600 | 7.28 | -0.03 | -19.26 | $< 10^{-75}$ |
| | 1800 | 7.40 | -0.02 | -16.95 | $< 10^{-59}$ |
| | 2000 | 7.51 | -0.01 | -7.30 | $< 10^{-12}$ |
| | 2200 | 7.61 | -0.01 | -10.19 | $< 10^{-23}$ |
| | 2400 | 7.70 | -0.01 | -10.67 | $< 10^{-25}$ |
| | 2600 | 7.78 | -0.01 | -8.65 | $< 10^{-16}$ |
| | 2800 | 7.85 | -0.01 | -10.30 | $< 10^{-23}$ |
| | 3000 | 7.92 | -0.01 | -7.98 | $< 10^{-14}$ |
| sLHP | 800 | 6.59 | 0.01 | 3.37 | $< 10^{-3}$ |
| | 1000 | 6.82 | -0.01 | -4.15 | $< 10^{-4}$ |
|  | 1200 | 7.00 | 0.00 | 1.86 | 0.06 |
| | 1400 | 7.16 | 0.00 | 2.71 | $< 10^{-2}$ |
|  | 1600 | 7.29 | -0.00 | -1.80 | 0.07 |
| | 1800 | 7.41 | -0.00 | -3.69 | $< 10^{-3}$ |
| | 2000 | 7.52 | 0.01 | 5.35 | $< 10^{-7}$ |
|  | 2200 | 7.61 | -0.00 | -0.21 | 0.83 |
| | 2400 | 7.70 | -0.00 | -3.17 | $< 10^{-2}$ |
|  | 2600 | 7.78 | -0.00 | -0.35 | 0.73 |
| | 2800 | 7.86 | -0.00 | -3.29 | $< 10^{-2}$ |
| | 3000 | 7.93 | -0.00 | -3.31 | $< 10^{-3}$ |
| GHP | 800 | 6.62 | -0.02 | -14.36 | $< 10^{-43}$ |
| | 1000 | 6.84 | -0.02 | -12.92 | $< 10^{-35}$ |
| | 1200 | 7.02 | -0.01 | -11.43 | $< 10^{-28}$ |
| | 1400 | 7.18 | -0.01 | -9.03 | $< 10^{-18}$ |
| | 1600 | 7.31 | -0.01 | -9.98 | $< 10^{-22}$ |
| | 1800 | 7.43 | -0.01 | -10.40 | $< 10^{-23}$ |
| | 2000 | 7.53 | -0.01 | -7.28 | $< 10^{-12}$ |
| | 2200 | 7.63 | -0.01 | -9.16 | $< 10^{-18}$ |
| | 2400 | 7.72 | -0.01 | -11.08 | $< 10^{-27}$ |
| | 2600 | 7.80 | -0.01 | -9.51 | $< 10^{-20}$ |
| | 2800 | 7.87 | -0.01 | -10.29 | $< 10^{-23}$ |
| | 3000 | 7.94 | -0.01 | -9.29 | $< 10^{-19}$ |

**Table S2.** Finite sampling bias corrections. *k* is the number of observed spontaneous replays after 10 000 s of spontaneous activity, *S*_*k*_ is the corresponding entropy of the assembly probability distribution based on replay count (Eq. 36) and *D*_*k*_ the diversity measure (Eq. 37). *Ŝ* is the finite sampling correction of the entropy and 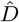 the corrected diversity calculated from the corrected entropy *Ŝ*.

| Stabilization rule | embedded assemblies | $k$ | $S$ | $\hat{S}_k$ | $D$ | $\hat{D}$ |
| --- | --- | --- | --- | --- | --- | --- |
| GHP | 800 | 92928 | 4.41 | 4.41 | 82.07 | 82.26 |
|  | 1000 | 79271 | 4.23 | 4.23 | 68.60 | 68.81 |
|  | 1200 | 49607 | 3.73 | 3.73 | 41.53 | 41.55 |
|  | 1400 | 27529 | 2.93 | 2.93 | 18.72 | 18.78 |
|  | 1600 | 9089 | 2.31 | 2.31 | 10.07 | 10.08 |
|  | 1800 | 1019 | 2.20 | 2.20 | 8.99 | 9.04 |
|  | 2000 | 18 | 1.80 | 1.97 | 6.04 | 7.16 |
|  | 2200 | 5 | 0.67 | 0.75 | 1.96 | 2.12 |
| sLHP | 800 | 82648 | 6.09 | 6.10 | 443.41 | 444.95 |
|  | 1000 | 69580 | 6.06 | 6.07 | 430.52 | 434.01 |
|  | 1200 | 60067 | 5.92 | 5.92 | 370.71 | 374.16 |
|  | 1400 | 52596 | 5.70 | 5.70 | 297.56 | 298.55 |
|  | 1600 | 36015 | 5.41 | 5.42 | 224.10 | 224.92 |
|  | 1800 | 22880 | 5.01 | 5.02 | 150.16 | 151.18 |
|  | 2000 | 12090 | 4.63 | 4.64 | 102.17 | 103.88 |
|  | 2200 | 8421 | 4.13 | 4.16 | 62.21 | 63.93 |
|  | 2400 | 4608 | 3.78 | 3.79 | 44.01 | 44.09 |
|  | 2600 | 2226 | 3.34 | 3.36 | 28.25 | 28.66 |
|  | 2800 | 1239 | 2.57 | 2.57 | 13.12 | 13.08 |
|  | 3000 | 368 | 2.20 | 2.21 | 9.04 | 9.11 |
| fLHP | 800 | 82560 | 6.31 | 6.32 | 551.36 | 552.83 |
|  | 1000 | 76871 | 6.31 | 6.32 | 549.28 | 552.82 |
|  | 1200 | 63618 | 6.18 | 6.19 | 485.22 | 485.41 |
|  | 1400 | 42930 | 6.05 | 6.06 | 423.90 | 427.33 |
|  | 1600 | 27075 | 5.78 | 5.79 | 324.00 | 326.28 |
|  | 1800 | 15879 | 5.34 | 5.35 | 209.32 | 210.60 |
|  | 2000 | 8518 | 5.01 | 5.03 | 149.39 | 153.36 |
|  | 2200 | 4433 | 4.58 | 4.59 | 97.22 | 98.05 |
|  | 2400 | 2114 | 3.97 | 4.03 | 52.86 | 56.31 |
|  | 2600 | 993 | 3.73 | 3.75 | 41.56 | 42.51 |
|  | 2800 | 480 | 2.77 | 2.77 | 15.89 | 16.00 |
|  | 3000 | 112 | 2.61 | 2.76 | 13.55 | 15.82 |

**Fig. S2.**
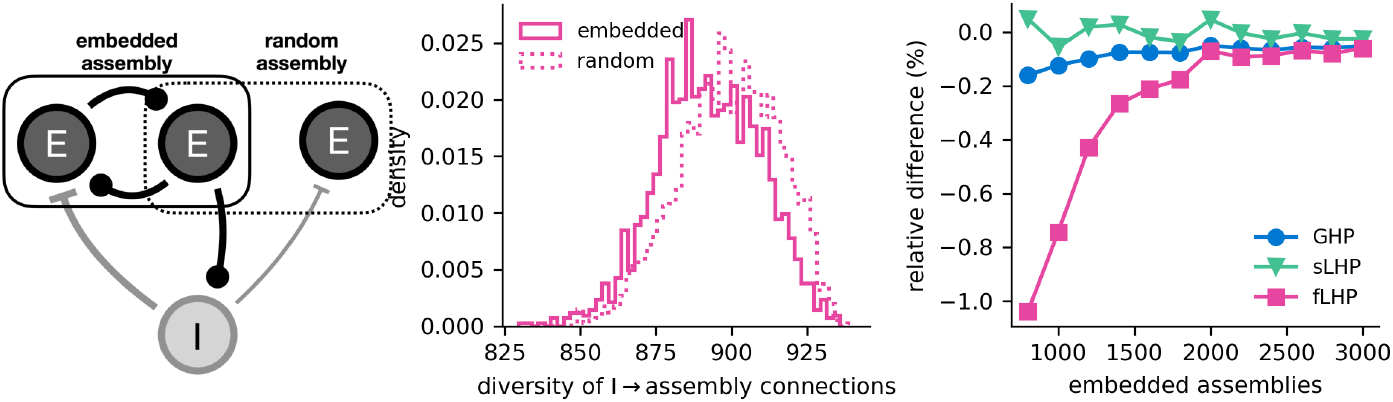
Specialization with embedded and random assemblies. **A:** An example connectivity scheme which can produce higher degree of inhibitory specialization on embedded assemblies, compared to random assemblies (shuffle control). The middle excitatory neuron (E_*M*_) excites the inhibitory neuron and the left excitatory neuron (E_*L*_). Activation of the middle excitatory neuron will thus lead to simultaneous firing of the inhibitory neuron and E_*L*_. Consequently, the inhibitory synapse from I onto E_*L*_ will strengthen more than the synapse from I onto the right excitatory neuron E_*R*_. **B:** Histogram of entropies calculated for each inhibitory neuron from its distribution of weights onto embedded assemblies and random assemblies (shuffle control - for each embedded assembly an assembly with the same number of neurons was assumed, but different composition). The histogram is shown for the fLHP rule where the change in the mean entropy is the largest (shown at 1000 embedded assemblies). Lower entropy with embedded assemblies means that inhibitory neurons are more specialized on embedded assemblies, as compared to the random shuffle control. **C:** Comparison of the relative change of the mean entropy from random assemblies to embedded assemblies. Statistical significance is evaluated in the Supplementary Table S1.

**Fig. S3.**
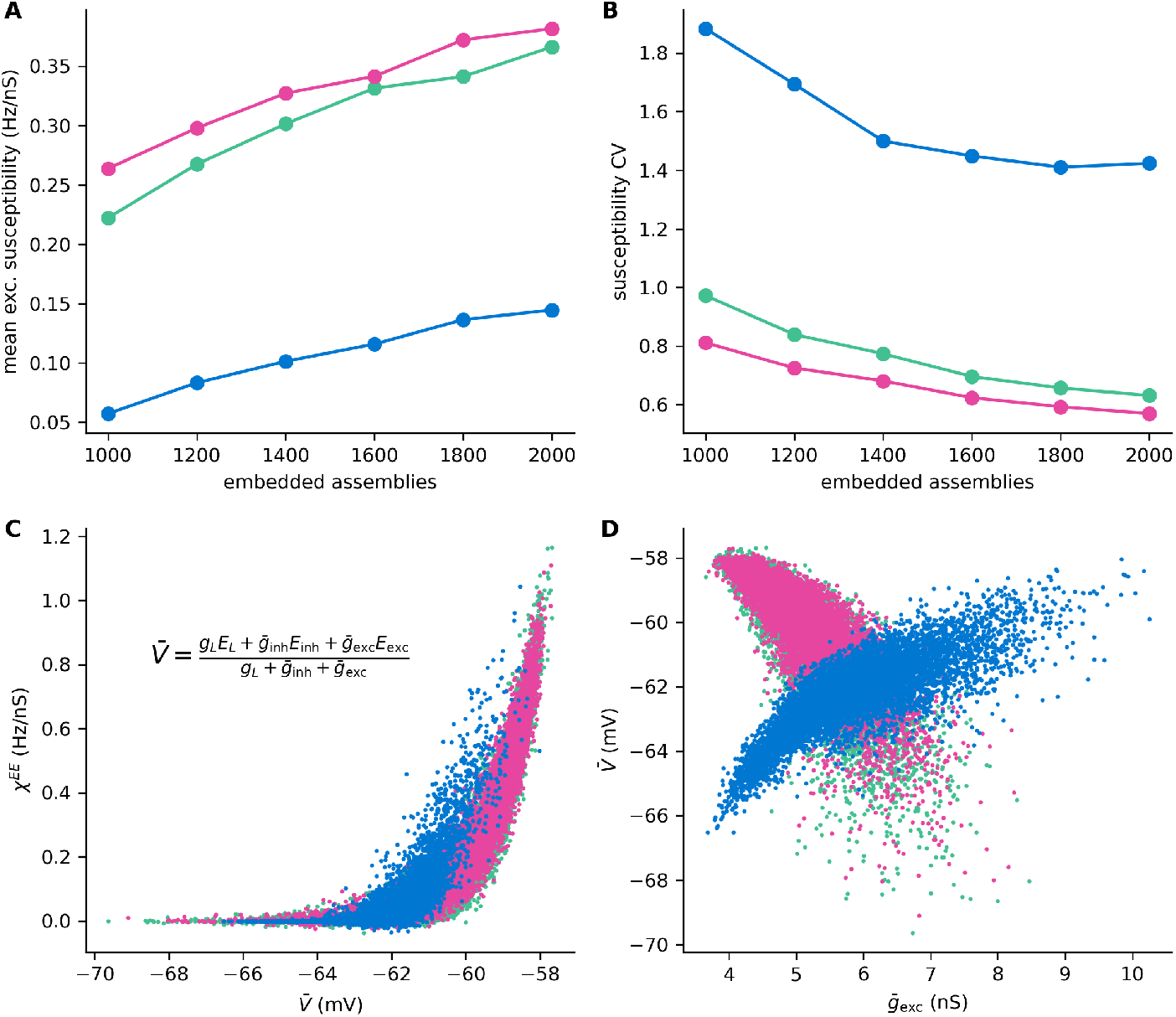
Sensitivity and its variability. **A:** Mean sensitivity to excitatory synaptic input for different numbers of embedded assemblies. **B:** Coefficient of variation the the sensitivity (standard deviation / mean). **C:** Dependence of the sensitivity on the resting potential, estimated as a weighted mean of reversal potentials. **D:** Dependence of the resting potential on the mean excitatory input.

**Fig. S4.**
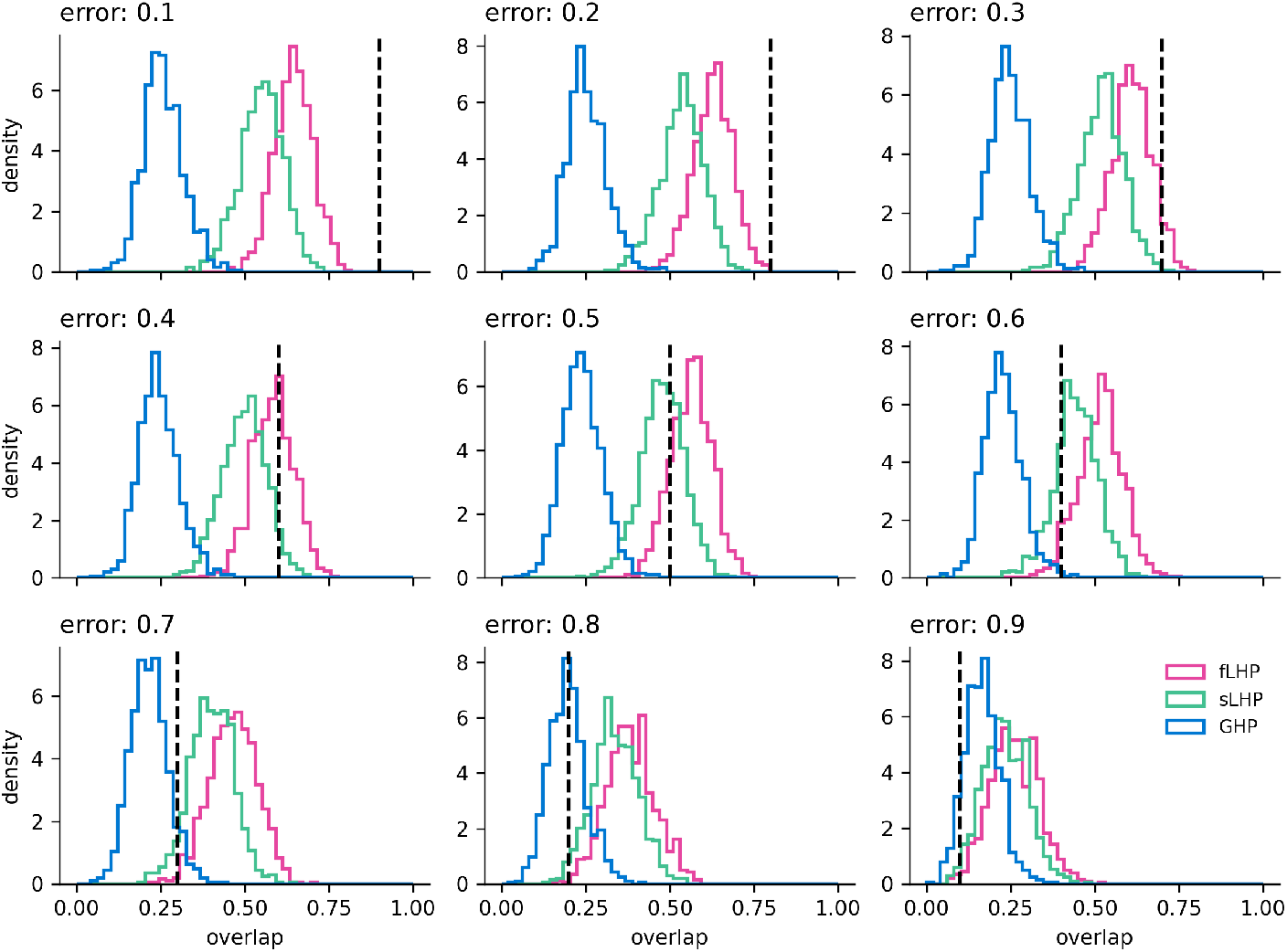
Pattern completion in linearized model. Histograms of overlaps of **W** · **c**_*m*_*u*(*e*) with *ξ*_*m*_*u*, where **c**_*m*_*u*(*e*) is the corrupted version of *ξ*_*m*_*u*. Any 1 in *ξ*_*m*_*u* is replaced with 0 with the error probability *e*. By default, the corrupted assembly **c**_*m*_*u*(*e*) has an overlap with *ξ*_*m*_*u* of (1 − *e*) (black dashed line).

## Notes

### Competing Interest Statement

The authors have declared no competing interest.

